# EZSolver: Template-free prediction of polar enzymatic mechanisms via bidirectional flow matching and search

**DOI:** 10.64898/2026.07.08.737313

**Authors:** Lun-Hsin Kuo, Jason Yang, Frances H. Arnold

## Abstract

Predicting enzymatic reaction mechanisms is critical for understanding enzyme function and for designing and discovering new enzymes. Current computational predictors rely on deterministic, rule-based dictionaries, which perform well on in-distribution tasks but fail to generalize to out-of-distribution (OOD) chemistry. To address this limitation, we present EZSolver, a template-free, generative framework for polar enzymatic mechanism prediction. Powered by a flow matching predictor (EZFlow) and navigated by an evaluator-guided bidirectional beam search, EZSolver learns the chemistry of electron redistribution instead of memorizing rigid templates. Evaluated across diverse enzyme classes, EZSolver achieves a 60.0% accuracy and an 84.6% chemical plausibility rate for full mechanism prediction of unseen polar enzymatic reactions. While rule-based models collapse without predefined templates, EZSolver successfully extrapolates chemical knowledge to infer uncatalogued pathways, as demonstrated during rigorous OOD benchmarking. By illuminating enzymatic chemical mechanisms, EZSolver helps pave the way for automated prediction of enzyme function and discovery and design of novel biocatalysts for sustainable chemistry.

**TOC Graphic:** 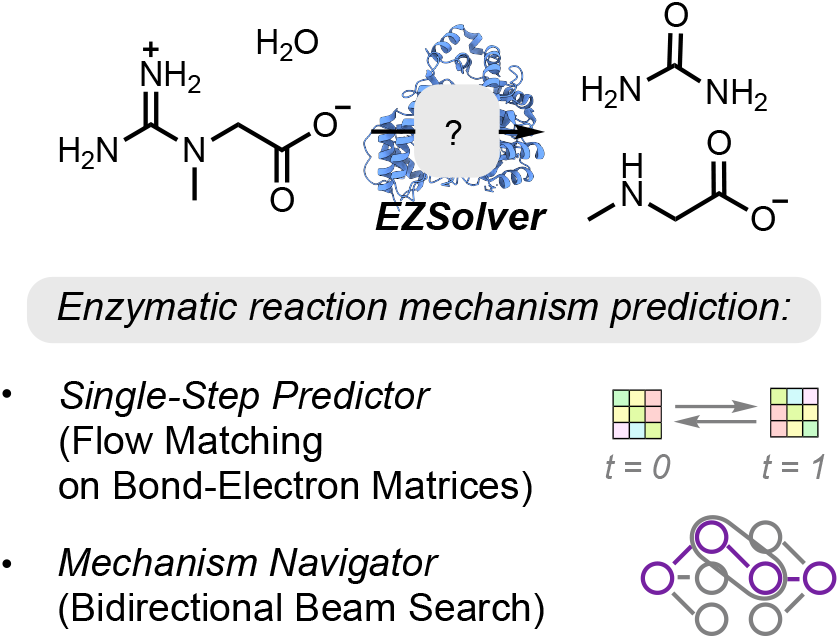

## INTRODUCTION

Nature’s remarkable protein catalysts—enzymes—hold the potential to transform how we make sustainable chemicals, fuels, and pharmaceuticals, carry out environmental remediation and crop protection, and process foods.^1–6^ Computational methods, including those powered by rapid advancements in artificial intelligence (AI) and machine learning (ML),^7–9^ are advancing our ability to discover or engineer enzymes and predict their functions. First, in enzyme discovery, deep learning frameworks^10–15^ are enabling us to mine sequence databases for new functions, enabling large-scale general functional annotation of uncharacterized proteins^10,16,17^ and discovery of specific enzymes, such as terpene synthases.^18,19^ Second, for enzyme optimization, machine learning-assisted directed evolution^20–23^ and zero-shot prediction models leverage evolutionary constraints and structural embeddings to navigate complex fitness landscapes. These models can significantly reduce the experimental burden required to enhance important enzyme properties such as catalytic efficiency, thermostability, and substrate specificity. Finally, generative AI models for sequence^24–27^ and structure-based design^28–30^ are unlocking *de novo* protein design capabilities, allowing researchers to computationally craft protein scaffolds, catalytic pockets, and functional enzymes from scratch.

Fundamentally, ML in biocatalysis aims to bridge the vast informational gap between the protein sequence-structure space and the chemical reaction space.^31–33^ The majority of current AI-driven biocatalysis tools have demonstrated efficient ways to encode enzyme sequences and structures. On the other hand, reactions are usually encoded solely as transformations from starting materials to products (Figure 1A).^34–36^ This approach hinders the model’s ability to understand key enzyme-substrate interactions, because the reaction mechanism is not represented. Practically, achieving reactions that are new to nature relies heavily on knowledge of reaction mechanisms, allowing scientists to discover the reaction promiscuity of proteins and expand catalytic scope toward novel chemical spaces.^37–41^ For AI models, neglecting valuable mechanistic information poses significant challenges in learning key protein-substrate interactions. Given that mainstream enzyme encoders (e.g., ESM) are trained primarily on evolutionary data that reflect existing enzymatic reactions, they intrinsically struggle to extrapolate to new-to-nature reactions. Consequently, incorporating mechanistic insights into generative models may enable the discovery and design of enzymes capable of catalyzing reactions beyond those discovered in nature.

**Figure 1.**
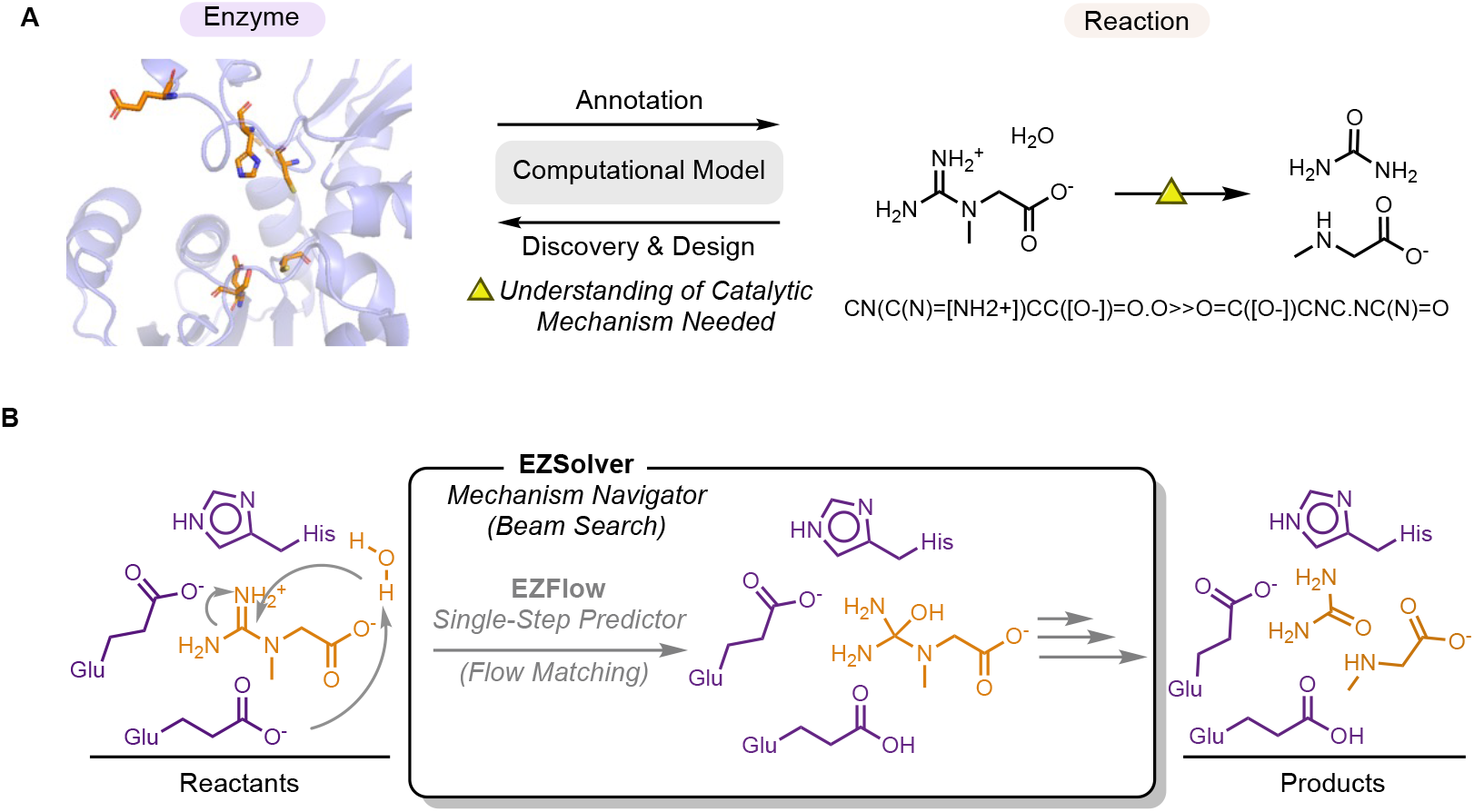
Overview of data-driven enzyme mechanism elucidation with EZSolver. (A) Neglecting mechanisms while training reaction encoder prevents models from learning real residue-substrate interactions. (B) EZSolver comprises a flow-matching-based single-step predictor (EZFlow) and a mechanism navigator.

Despite the critical need for mechanistic insights, mechanistic data for enzymatic reactions are exceedingly limited, and predicting enzymatic mechanisms remains a formidable computational challenge. Recent advancements, such as EZMechanism^42^ and MechFind,^43^ have made progress in enzymatic mechanism retrieval and data curation.^44^ By utilizing algorithms like heuristic graph search or mixed-integer linear programming (MILP), these models successfully retrieve discrete reaction rules that sum up to the overall chemical transformation.^43^ However, these methods rely fundamentally on fixed dictionaries of predefined elementary rules extracted from existing databases, which strictly limits their ability to tackle out-of-distribution tasks. There has been separate progress for predicting the outcomes of organic reactions. These include transformer-based models (e.g., T5Chem and Chemformer),^45–47^ graph-based models (e.g., WLDN, LocalRetro, and Graph2SMILES),^48–50^ and flow matching-based models like FlowER.^51^ However, these models were primarily evaluated in product prediction and lack the chemical intuition required to reveal accurate mechanisms. To address this limitation, Miller et al.^52^ developed PMechRP, a hybrid framework powered by a Chemformer ensemble that incorporates a Siamese neural network to evaluate proposed elementary steps. Yet, when tested on enzymatic reactions, its base model (Chemformer) suffers from severe model hallucinations. To the best of our knowledge, a template-free, generative architecture capable of elucidating enzymatic reaction mechanisms is lacking.

Here, we present EZSolver, a generative framework designed specifically to navigate multi-step polar enzymatic mechanisms. Our approach features a powerful two-component architecture: a flow-matching-based single-step predictor, EZFlow, and an evaluator-guided mechanism inference navigator, EZSolver (Figure 1B). EZFlow builds off the architecture of FlowER,^51^ preventing hallucinations by modeling chemical transformations through continuous electron redistribution within a Bond-Electron (BE) matrix. In addition, we use both forward and inverse flow to predict reversible reactions, which are highly prevalent in enzymes. Finally, we assemble these elementary steps into biologically relevant pathways through a bidirectional beam search algorithm directed by a customizable dual-evaluator system (EZSolver). Ultimately, EZSolver outputs multiple feasible mechanisms and ranks them based on step length and an internal chemical scoring metric. In our out-of-distribution evaluation, EZSolver achieved 60.0% accuracy on full mechanism prediction, and 84.6% of the reactions were proposed with chemically plausible mechanisms. Benchmarking on the out-of-distribution task from EZMechanism^42^ yielded a 52.7% accuracy for EZSolver, substantially outperforming existing rule-based models (23.6% for EZMechanism and 18.8% for MechFind). This exceptional generalizability and robust chemical intuition highlight the distinct advantages of generative architectures over deterministic rule-based models. By synergizing chemistry-informed flow matching with structurally guided search algorithms, EZSolver effectively conquers the combinatorial explosion of reaction space, offering a robust prediction engine for polar enzymatic reactions. The code for training and using EZFlow and EZSolver is publicly available at https://github.com/fhalab/EZSolver.

## METHODS

### Datasets and splitting

We focused on predicting mechanisms for polar and non-metallic reactions, a major category in enzyme catalysis (acidbase catalysis).^53^ After curation and filtration of the M-CSA database,^54^ we obtained a dataset comprising approximately 1,200 elementary steps with accurate atom mapping. We acknowledge that curating the exact proton transfers remains fundamentally challenging. While our automated heavy-atom and explicit hydrogen mapping protocol achieves high fidelity for the reaction skeleton, the subtle ambiguity in assigning exact protonation states to spectator residues represents an inherent limitation in data curation. A subset of 186 elementary steps, comprising 65 full reaction mechanisms, was selected for testing model performance in most results presented in this study. This testing dataset was used to examine the prediction accuracy of the models in both elementary step predictions and full mechanism predictions. No elementary step in the testing set shares an identical SMILES string with the training data. Further analysis of the Reaction Tanimoto Similarity between elementary steps from the training and testing sets revealed that roughly 13% of the testing data exhibited a 100% maximum reaction similarity compared to the training data (Figure S1). Because these elementary steps are almost exclusively proton transfers between amino acid residues, which are shared across nearly all enzymes, we can rule out the possibility of data leakage (i.e., the model achieving high testing accuracy by simply memorizing the training data). Additional details on dataset curation are presented in the Supporting Information.

### EZFlow: elementary step prediction with flow matching

Driven by flow matching, EZFlow predicts elementary steps by modeling electron redistribution within a Bond-Electron (BE) matrix, an adjacency matrix where the rows and columns represent individual atoms, and the matrix elements denote the number of bonding or non-bonding electrons. Inheriting the organic chemistry knowledge from its predecessor, FlowER,^51^ EZFlow successfully learns the functionalities of amino acid residues from limited enzymatic data and incorporates bidirectional flow matching to predict reversible reactions.

#### Finetuning

The single-step predictor, EZFlow, is obtained by finetuning FlowER, using the same attention-based architecture on the BE Matrix of chemical compounds. Finetuning initiates from the pretrained weights of FlowER to transfer chemical knowledge from organic reactions. We deliberately substitute the element Europium (Eu), which is nearly absent in the FlowER training data, with the character (‘*’) to represent arbitrary R-groups within organic structures. Finetuning of FlowER was conducted with a learning rate of 1 × 10^−5^, a dropout rate of 0.1, 2,200 warmup steps under a Noam learning rate scheduler, and a training Gaussian noise standard deviation (σ) of 0.05. The optimal checkpoint was selected based on inference accuracy, evaluated by converting the predicted BE matrix back into SMILES strings for direct comparison with the ground-truth product SMILES. The sample size *(N)* represents the number of independent trajectories generated from initial prior distribution during flow matching inference. In training, the validation accuracy was evaluated using a fixed sample size of *N = 16*. In evaluation, sample size varies, and the corresponding impacts are discussed in the *Results and Discussion* section. Hyperparameters for EZFlow model training are found in Table SI of Supporting Information.

#### Inverse flow

To predict the reversible elementary steps prevalent in enzymatic reactions, we conceptualize the flow matching trajectory as the thermodynamic vector field (Figure 2A). While standard inference integrates from reactant state (t = 0) to product state (t = 1), the inverse flow protocol reverses the time integration in flow matching. During inference, predictions of standard and inverse flow are combined to explore the entire thermodynamic landscape (Figure 2B).

**Figure 2.**
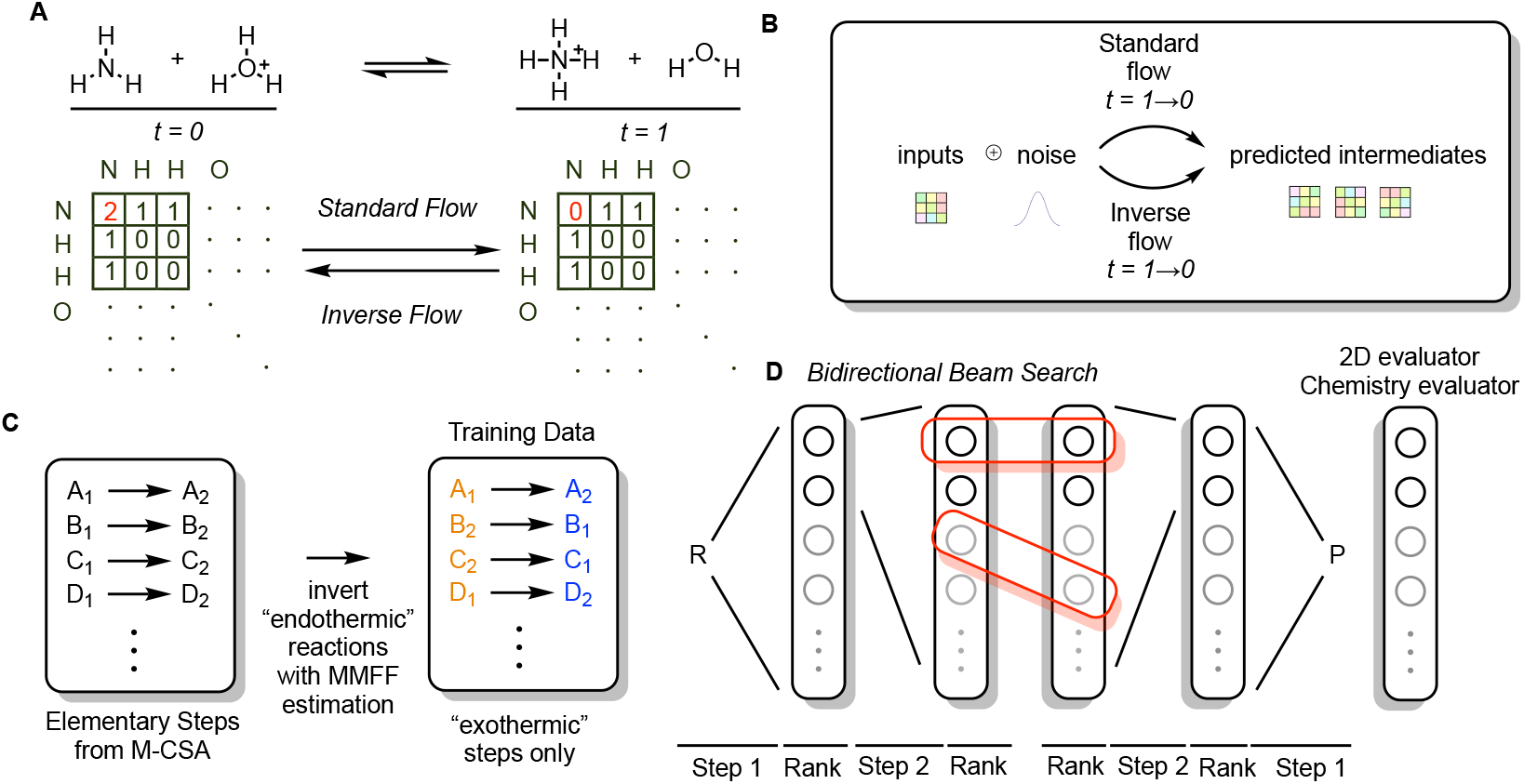
Key architectural components of EZFlow and EZSolver. (A) Inverse flow allows EZFlow to infer reversible elementary steps, which are prevalent in enzymatic reactions. (B) EZFlow takes the union set of predictions from standard flow and inverse flow as the output in each round of inference. (C) Processing reactions with MMFF estimation prevents contradictory signals during training and improves model performance. (C) Architecture of the mechanism navigator. Applying bidirectional beam search with an expert-designed evaluator allows the mechanistic inference process to focus on relevant chemical space, ultimately achieving a high success rate in identifying reaction mechanism between given reactants and products.

#### MMFF

To evaluate the reaction energy profile, Merck Molecular Force Field (MMFF) in RDKit was utilized to estimate the potential energy difference between reactants and products. To prevent contradictory gradient signals during EZFlow training, the dataset was aligned into an “exothermic” set by reversing all the endothermic reactions (Figure 2C).

#### Focal Weight

Because a typical elementary step involves electronic reorganization among only a few reactive centers, the vast majority of elements in a BE matrix remain static. This extreme sparsity dilutes the gradient signal for actual chemical transformations, making it difficult for the model to capture localized reactivity. To overcome this bottleneck, a focal penalty factor, W_ij_, is introduced into the loss function to amplify gradient signals on chemically reacting atoms while suppressing the static background. Chemically reactive atoms were identified by comparing the BE matrices of the current states to the final states. The focal weight is dynamically updated at every round of inference. The training objective is defined in Equation 1, where M_ij_ denotes the atom mask matrix, W_ij_ represents the focal weight, v_t,ij_ corresponds to the predicted vector field, and u_t,ij_ is the ground-truth vector field. The optimal focal weight ratio for W_ij_ was found to be 4:1 for changed versus unchanged elements in the BE matrix.

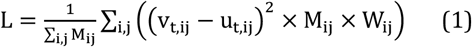

#### Noise Standard Deviation (σ)

During inference, we explored whether elevating the standard deviation (σ) can encourage the model to explore a wider chemical space, thereby enhancing accuracy. While excessively high σ values induce model hallucinations, the optimal inference noise was empirically determined to be (σ = 0.08).

### EZSolver: full mechanism prediction with bidirectional beam search

Given reactants and products, EZSolver performs an exhaustive bidirectional beam search, utilizing EZFlow as the generative single-step predictor, to construct multiple plausible reaction mechanisms. These generated pathways are subsequently ranked based on their total step length and internal scores derived from customized evaluators.

#### Bidirectional beam search

Multi-step mechanism prediction is notoriously plagued by error accumulation, where a single incorrect elementary step derails the entire subsequent pathway. To overcome this, we developed a bidirectional beam search algorithm that performs inference simultaneously from both the reactant and product states (Figure 2D). Importantly, unlike greedy search algorithms that terminate immediately upon finding a single path, our bidirectional beam search is designed to grow continuously until reaching a predefined maximum depth (or until all beams naturally terminate). Once completed, the algorithm finds intermediate nodes with identical chemical structures from both beams and traces back to their corresponding roots (reactants and products) to construct the full mechanism. This exhaustive exploration ensures that the algorithm does not merely identify the shortest path, but comprehensively maps out multiple coexisting mechanistic proposals within the chemical landscape. To effectively navigate this vast search space and pinpoint the most plausible and accurate mechanisms, we implemented a built-in dual-evaluator system consisting of a *2D Navigator* and a *Chemistry Evaluator*, which are used to prune the search trees while maintaining a predefined beam width until maximum beam depth. The detailed algorithm for EZSolver is provided in the Supporting Information, and the hyperparameters used during inference are listed in Table S1.

#### 2D Navigator

By calculating the change in the BE matrix between the intermediate and final states, the 2D evaluator identifies atoms experiencing reactions and applies a weighting matrix to reward steps aligned with the overall reaction. Crucially, this filter eliminates chemically accurate but irrelevant intermediates early on, effectively constraining the beam search to focus solely on the chemically relevant path.

#### Chemistry Evaluator

The Chemistry Evaluator acts as a rigorous chemical sanity check. This module penalizes the formation of highly reactive, thermodynamically improbable intermediates, or chemical hallucinations. By integrating these evaluators to rank candidates, the algorithm filters out high-probability but chemically absurd shortcuts, ensuring only the top-k most chemically sound candidates survive to the next iteration.

#### Evaluation

After completing search, we ranked the retrieved mechanisms firstly based on the length of mechanisms and followed by the sum of scores from the evaluators into the top “k” proposals, where k = 1, 5. Every proposed pathway was manually inspected by experts and classified as ‘accurate’, ‘chemically plausible’, or ‘retrieved’ according to mechanistic criteria.

## RESULTS AND DISCUSSION

### Performance of elementary step prediction with EZFlow

Initial testing of the standard FlowER architecture on the enzymatic elementary reactions yielded a mere 3.3% accuracy. Common elementary reactions that the model failed to predict included proton transfers that violate pKa trends, concerted general acid/base mechanisms, and the formation of zwitterions. Due to the hydrophilic environment and complicated hydrogen bonding network in the enzyme pocket, the thermodynamic landscape of intermediates deviates dramatically from that in organic solvents. To address this, our attempt of directly *finetuning* FlowER on the enzymatic training set converged to a maximum 35.7% top-200 accuracy. After detailed investigation, we found that this problem stems from the inherent nature of biochemical pathways, the pervasive presence of reversible reaction pairs within the training data (e.g., reversible proton transfers and substrate residue interactions). Since FlowER operates fundamentally as a single-directional reaction predictor, it struggles to learn from the contradictory objective vector fields generated by these reversible reaction pairs. Our key innovation to resolve this limitation is allowing for both forward and reversed time integration during inference. This *inverse flow* approach enables the prediction of elementary steps that are oppositely aligned with the training set (see Supporting Information for detailed discussion). Combining the prediction from standard and inverse flow boosts the accuracy to 50.1%. By processing the training dataset to “exothermically” aligned reactions through MMFF estimation (*MMFF*), the vector field contradiction was further mitigated in training, resulting in a 60.3% prediction accuracy. Additionally, using the optimal hyperparameter of the weight matrix (*Focal weight*) boosted performance to 66.1% accuracy. Lastly, during the inference stage, we deliberately *increased the standard deviation (σ)* of initial Gaussian noise distribution to explore a broader chemical space, which ultimately boosted the elementary step prediction accuracy to 72.4% (Figure 3A). From the ablation study, we conclude that the flow matching ODE could be reconceptualized not merely as a generative trajectory, but as a bidirectional thermodynamic vector. Training on the exothermically aligned dataset and incorporating inverse flow during inference allow EZFlow to explore the entire thermodynamic landscape, resulting in optimal accuracy on the unseen testing set.

**Figure 3.**
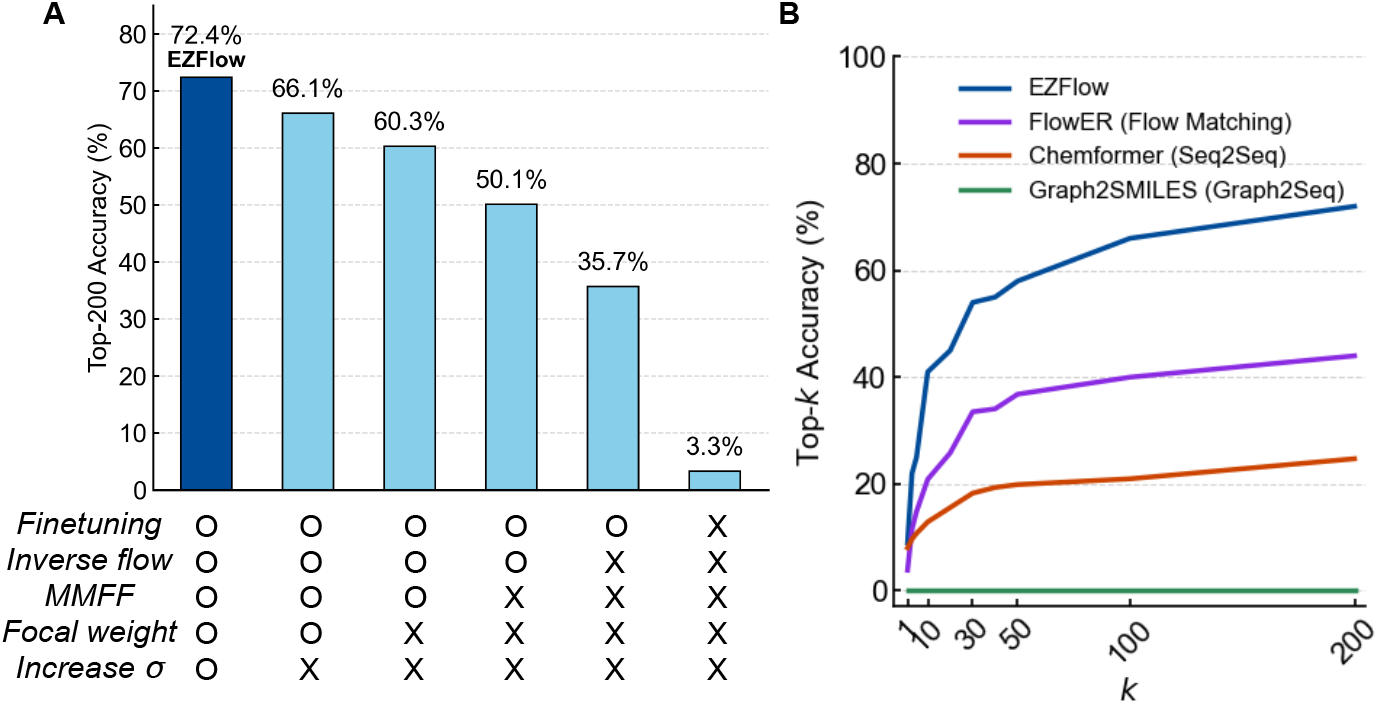
Bidirectional flow matching framework of EZFlow showing advantages over FlowER and deterministic models. (A) Ablation study of EZFlow, showing how each method design choice (elaborated in *Methods*) improves performance, showing percentage accuracy for when an exact match to the correct elementary step can be found within the top 200 proposals. (B) Benchmark against fine-tuned FlowER, Chemformer, and Graph2SMILES, with all baseline finetuning executed in their respective original protocols. With the prior knowledge from the ablation of EZFlow, we adapted focal weight and increased *σ* in finetuning and evaluating FlowER.

### EZFlow suggests reasonable elementary steps despite complex molecular inputs

To contextualize the performance of EZFlow, we benchmarked the method against state-of-the-art models, including Chemformer^47^ (a transformer-based sequence-to-sequence model) and Graph2SMILES^50^ (a graph-to-sequence model). To ensure a rigorous and fair evaluation, we fine-tuned all pretrained baseline models exclusively on the identical enzymatic training set. The comparative top-k predictive performance of these models is presented in Figure 3B, where the top-200 accuracies of EZFlow, FlowER, Chemformer, and Graph2SMILES are 72%, 35%, 24%, and 0%, respectively.

The increase in chemical complexity between standard organic synthesis and enzymatic catalysis may explain why predicting enzymatic elementary steps is particularly difficult. While models trained on the USPTO dataset encounter comparable reaction complexities in terms of average heavy atom counts, the nature of the spectator molecules differs drastically from that in enzymatic reactions. In organic datasets, spectators are predominantly solvents or counter-ions (e.g., THF, K^+^, Cl^-^), allowing models to readily ignore them as background noise. Conversely, spectator molecules in enzymatic reactions are amino acid residues within the catalytic pocket. These residues interact with the substrate across different elementary steps, meaning models cannot treat them as benign noise; they must rationally identify the correct interacting residue. This massively expands the chemically valid elementary step space and makes it extremely challenging for deterministic models like Graph2SMILES and Chemformer to predict the correct elementary step. This challenge is reflected in Chemformer’s performance (Figure 3B), which achieved a high Top-1 Tanimoto similarity (76%) but a catastrophic Top-1 exact-match accuracy (8%). The massive discrepancy highlights a fundamental limitation of deterministic sequence-to-sequence architectures in enzymatic reaction prediction.

An example of Aralkylamine N-acetyltransferase (AANAT)-catalyzed acetylation of serotonin is illustrated in Figure 4A. The first elementary step involves a histidine-assisted nucleophilic attack from serotonin to acetyl-CoA (indicated by red arrows). This crucial step was successfully predicted by EZFlow but failed entirely with Chemformer.^47^ Notably, while 97% of the intermediates predicted by Chemformer were chemically valid, none of them showed the conservation of atoms (Figure 4A). This catastrophic failure can be attributed to the structural complexity of acetyl-CoA and, most importantly, the vastly enlarged scope of chemically valid steps introduced by the coexistence of reactive residues like serine, histidine, and tyrosine in the active site. In contrast, EZFlow navigated this complexity, predicting the true intermediate. By sampling the BE matrix from diverse initial noise distributions, EZFlow mitigates the deterministic bias inherent in sequence-to-sequence models and enforces atom conservation at the same time. Notably, other chemically valid elementary steps were also predicted by EZFlow. The blue and green arrows highlight two selected examples of valid elementary step predictions, including alternative proton transfers and a histidine-assisted nucleophilic attack from a water molecule to acetyl-CoA. While this multiplicity initially increases the difficulty of the search process, it is the key to ensuring a comprehensive exploration of the enzymatic reaction space, which is ultimately managed and navigated by EZSolver’s evaluators.

**Figure 4.**
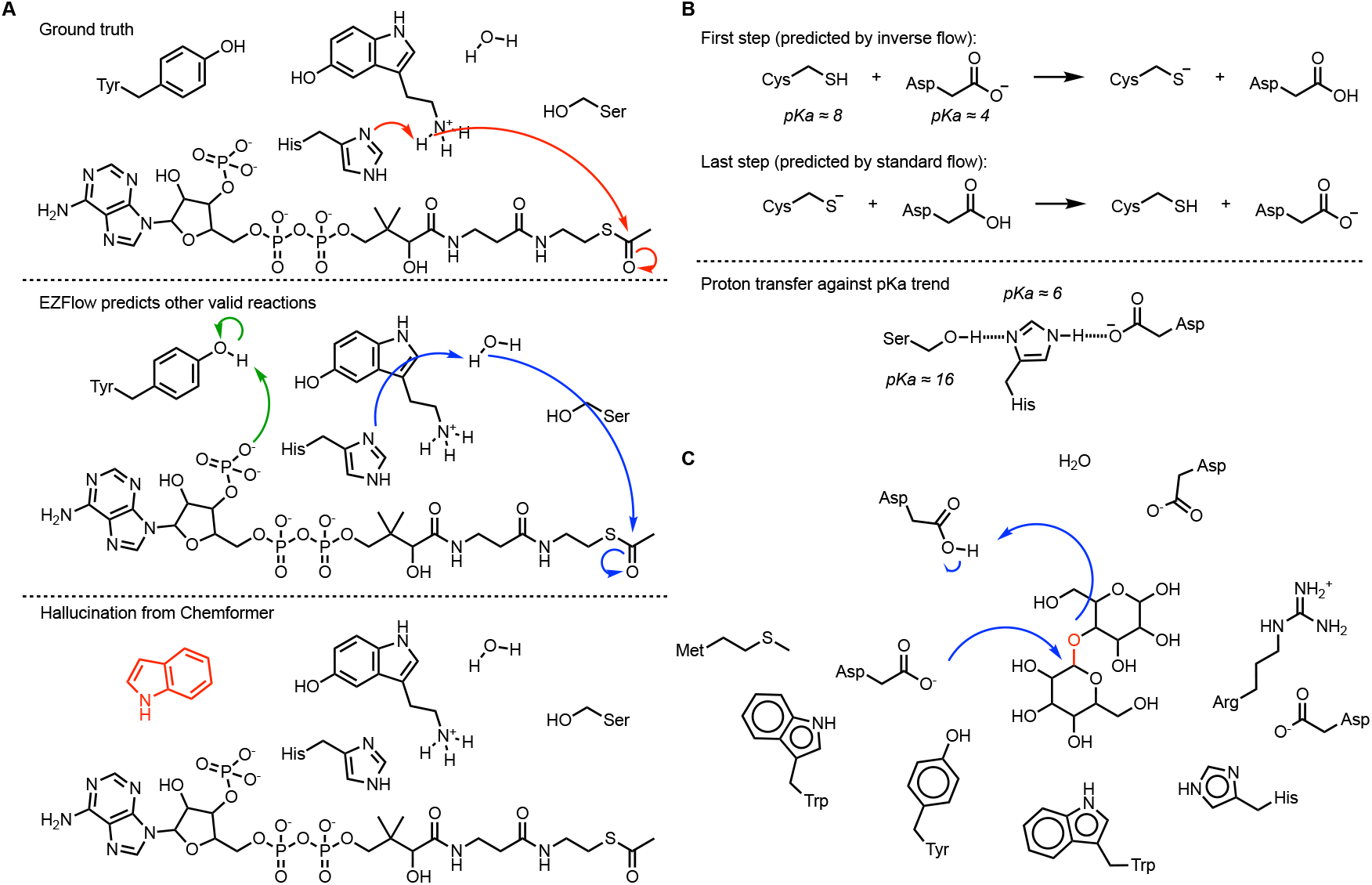
Case studies highlight the advantages of EZFlow in preventing hallucinations, predicting reversible reactions, and handling complex reaction inputs. (A) An example of serotonin acetylation reflects the generalizability of EZFlow and the limitations of deterministic models like Chemformer. (B) The coexistence of reversible elementary steps in glutamate racemization showcases the importance of combining inverse flow in solving enzymatic reaction mechanisms. (C) A maltose hydrolysis example demonstrates the utility of focal weighting, allowing EZFlow to handle complex inputs.

Figure 4B illustrates the importance of inverse flow. After training on predominantly exothermic reaction data, standard flow matching vectors intrinsically acquire downhill thermodynamic characteristics. This bias is evident in the epimerization catalyzed by glutamate racemase, which involves a thermodynamically uphill proton transfer between aspartate and cysteine in the initial step, followed by the reverse (downhill) reaction in the final step. Constrained by its exothermic training bias, the fine-tuned FlowER model failed to predict the initial uphill proton transfer. In contrast, EZFlow successfully predicted the uphill initial step through inverse flow, while accurately resolving the downhill final step via standard flow. Furthermore, this capability extends to fundamental enzymatic functionalities, such as the activation of the catalytic triad to form a nucleophilic serine intermediate—a process that inherently contradicts aqueous pKa trends. By strategically combining standard and inverse flow, EZFlow overcomes these thermodynamic problems, enabling the accurate prediction of elementary steps across the entire thermodynamic spectrum.

EZFlow also has excellent capacity to manage highly complicated tasks. This is illustrated by the initial step of maltose hydrolysis, which involves 12 compounds in the starting state (Figure 4C). The fine-tuned FlowER model predictably failed to identify the correct elementary step. In contrast, EZFlow successfully overcame this chemical noise, correctly pinpointing the exact roles of specific residues, selecting an aspartate as the nucleophile and a protonated aspartic acid as the proton donor to achieve the precise fragmentation of maltose. This successful prediction directly highlights the critical contribution of the focal penalty weight (W_ij_). By suppressing the influence of the numerous spectator residues, the focal penalty algorithm forces the generative flow to concentrate exclusively on the localized active centers. This enables EZFlow to resolve complex multi-residue systems.

### Performance of full mechanism prediction with EZSolver

Enzymatic reactions exhibit mechanistic multiplicity. This is reflected in various proposals for each reaction in the M-CSA database, where those plausible pathways may coexist under physiological conditions. For instance, the hydrolysis of a glycosidic bond may be rationalized by an S_N_2 mechanism involving specific residues or a stepwise path via an oxocarbenium intermediate. Consequently, evaluating the mechanism proposed by the model purely on a binary “correct/incorrect” criterion fails to capture its true chemical intuition. To provide a comprehensive analysis, we individually screened and categorized the prediction results into three distinct levels (examples shown in Figure 5A). The “Accurate” level requires the generation of the correct intermediates and the involvement of accurate substrate-residue interactions, while relaxing the constraints on the exact transient positions of protons due to the inherent ambiguity of protonation states within the hydrogen-bonding network. The “Chemically Plausible” level denotes that the model generates an alternative mechanism that follows rigid physical and chemical rules. For example, instead of direct hydrolysis, a predicted serine-catalyzed hydrolysis mechanism is categorized as Chemically Plausible (Figure 5A). Finally, the “Retrieved” level represents scenarios where the system successfully discovers a complete mechanism connecting the initial reactants to the final products but disobeys common chemical rules, such as by generating extremely unstable intermediates.

**Figure 5.**
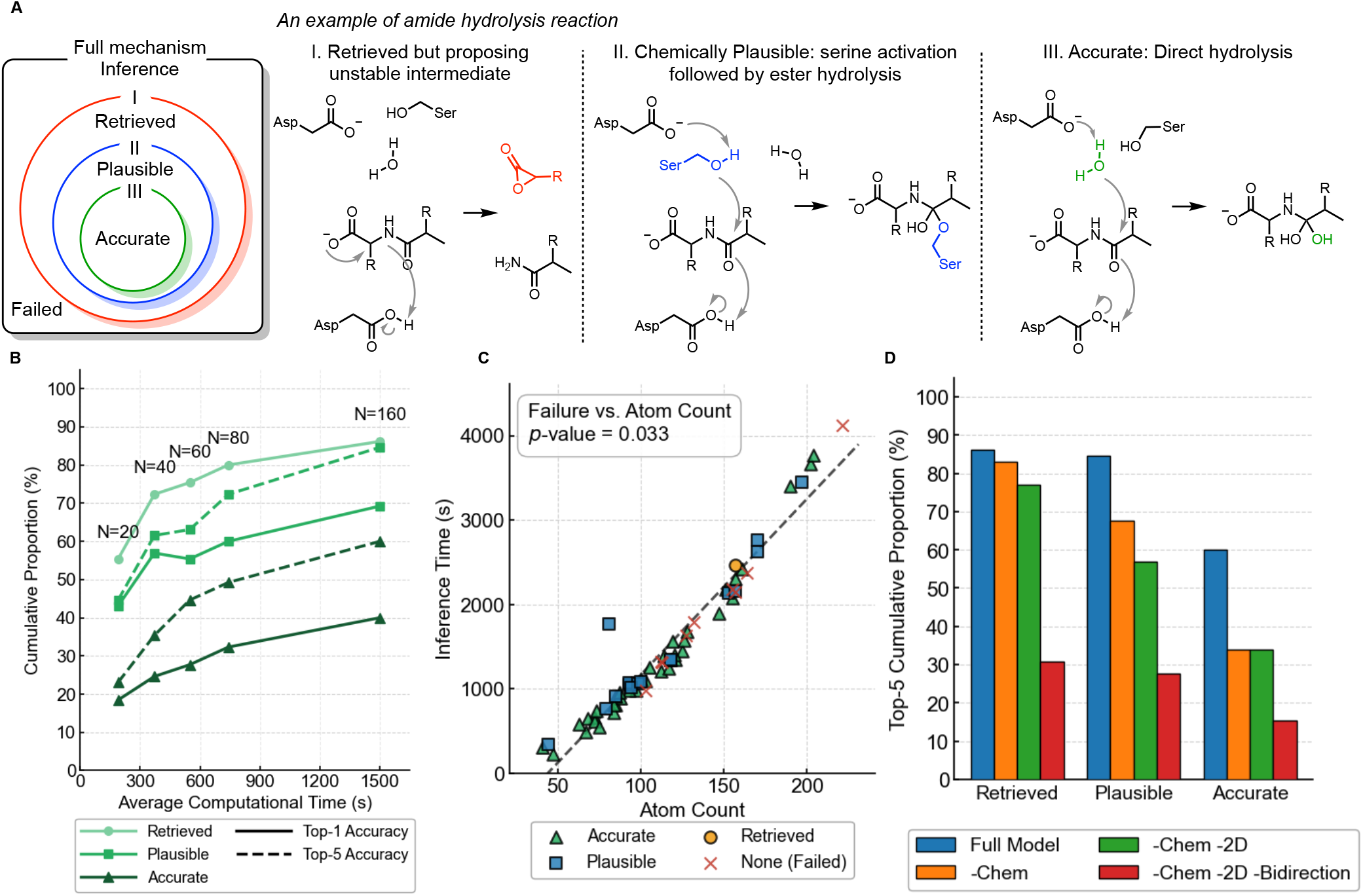
EZSolver mitigates error accumulation during inference and achieves high full-mechanism prediction accuracy. (A) Due to the mechanistic multiplicity of enzymatic reactions, we define mechanism proposals at three levels: Retrieved, Chemically Plausible, and Accurate. The existence of the ‘Chemically Plausible’ category reflects the model’s ability to generalize learned chemical knowledge into an enzymatic context, a capability absent in rule-based models. (B) Computational time versus the proportion of each accuracy level. (C) Inference time for each reaction shows a linear correlation with atom count. While failure cases show a p-value of 0.033 relative to atom count, they are distributed across a broad spectrum of molecular sizes. This indicates that EZSolver is not solely hindered by the atom count of reactions. (D) Ablation study of the mechanism navigator shows the importance of each component. -Chem: removing chemistry evaluator. -2D: removing 2D navigator. -Bidirection: conducting single directional beam search from the reactants only.

Our bidirectional mechanism searching model, EZSolver, shows strong performance when evaluated using these metrics. Figure 5B shows the analysis of computational cost and predictive accuracy through modulating the sampling size (N) in the flow matching process. Increasing the sample size allows the model to extensively explore the space of possible elementary steps. Under the navigation of the two evaluators, EZSolver robustly secures perfect mechanisms within the top-5 candidates, achieving a 60.0% success rate for Accurate and an 84.6% rate for Chemically Plausible mechanisms. Notably, nearly all retrieved mechanisms within the top-5 candidates are chemically plausible when the *N* is scaled to 160. Instead of being confined by predefined reaction templates like traditional rule-based algorithms, our generative model explores the vast chemical space without template restrictions and proposes a diverse ensemble of chemically sound mechanisms.

Inherently more complex reactions (with more atoms) tend to require greater computational cost for mechanism prediction and are generally more difficult for EZSolver to predict accurately. In Figure 5C, the inference time exhibits a linear correlation with the input atom count (dimension of BE matrix in flow matching). Although molecule size has a statistically significant effect on prediction failure rate (p value = 0.033), a closer examination of the distribution reveals that failures are not exclusively confined to large and complicated systems. Instead, unsuccessful predictions are distributed across a broad spectrum of molecular sizes. This distribution pattern suggests that while the sheer number of heavy atoms marginally increases the combinatorial difficulty, the primary bottleneck for prediction success lies in the intrinsic chemical complexity of the transformation, such as intricate proton-shuttling networks or highly unusual reaction trajectories, rather than merely the size of the input.

To systematically understand the contribution of each method design component, we performed the ablation study shown in Figure 5D. The essentiality of the *Chemistry Evaluator* is apparent upon its removal. When the chemical sanity checks are disabled, the overall retrieval rate remains predictably high (83.1%), as the model is still capable of structurally connecting reactants to products. However, without strict chemical penalties, the search algorithm begins to exploit high probability but chemically ridiculous shortcuts. Common mistakes include proposing highly strained cyclic intermediates or skipping tetrahedral intermediates during substitution on carbonyl derivatives. This leads to a significant divergence from the ground truth in over half of the retrieved reactions (Plausible: 67.7%, Accurate: 33.8%). Further removing the *2D Navigator* deprives the search algorithm of its topological focus. Without the structural guidance derived from the Bond-Electron matrix to steer the pathway progressively toward the target product, the network wanders aimlessly in the vast chemical space, causing the three performance metrics to decline simultaneously (Retrieved: 76.9%, Plausible: 56.9%, Accurate: 33.8%). Finally, degrading the system to a unidirectional baseline model—by removing the *bidirectional search* along with both evaluators—results in the biggest performance drop across the board (Retrieved: 30.7%, Plausible: 27.7%, Accurate: 15.4%). This catastrophic degradation underscores that the astronomically vast combinatorial space of enzymatic elementary steps poses a significant search challenge that cannot be overcome by naive unidirectional inference. Interestingly, even in this fully ablated baseline state, the Plausible rate remains strikingly close to the Retrieval rate, and half of the retrieved mechanisms propose key intermediates and correct substrate-residue interactions. This demonstrates that the foundational pre-training of the single-step predictor, EZflow, was highly successful in inherently embedding a strong baseline of chemical intuition into the model’s neural weights, ensuring that even its unguided proposals remain largely grounded in chemical reality.

Figure 6A illustrates a successful ‘accurate’ case involving the aldol reaction catalyzed by 2-dehydro-3-deoxy-phosphogluconate aldolase. Without relying on predefined reaction templates, EZSolver correctly identified lysine as the nucleophilic catalyst, initiating the formation of a critical enamine intermediate. The subsequent nucleophilic attack on the aldehyde followed by the hydrolysis of the iminium species successfully yielded the aldol products. This example underscores the competence of the bidirectional search algorithm in navigating long-range mechanistic pathways. The combined guidance of the 2D Navigator and Chemistry Evaluator efficiently pruned the vast chemical search space, while the foundational chemical intuition embedded during EZFlow’s training allowed the model to correctly assign catalytic functionalities to specific activesite residues. Figure 6B presents two distinct mechanism proposals generated by EZSolver for a desulfination reaction. In these two scenarios, the active-site cysteine plays different catalytic roles: in one pathway, it acts as a general acid triggering an electrophilic aromatic substitution; in the other, it acts as a nucleophile activating the sulfinate group for subsequent substitution. These two mechanisms were actually the subject of a long-standing debate in the scientific community.^55^ While early crystallographic evidence favored the latter nucleophilic mechanism,^56^ subsequent DFT calculations ultimately confirmed the former EAS pathway as the more plausible alternative.^57^ Notably, this is the only desulfination reaction in the entire M-CSA database, meaning EZSolver had no prior exposure to this reaction class during training. The recovery of both mechanisms without any reliance on crystallographic data or DFT calculations highlights its ability to extrapolate to unseen reaction types and propose chemically insightful pathways for further analysis.

**Figure 6.**
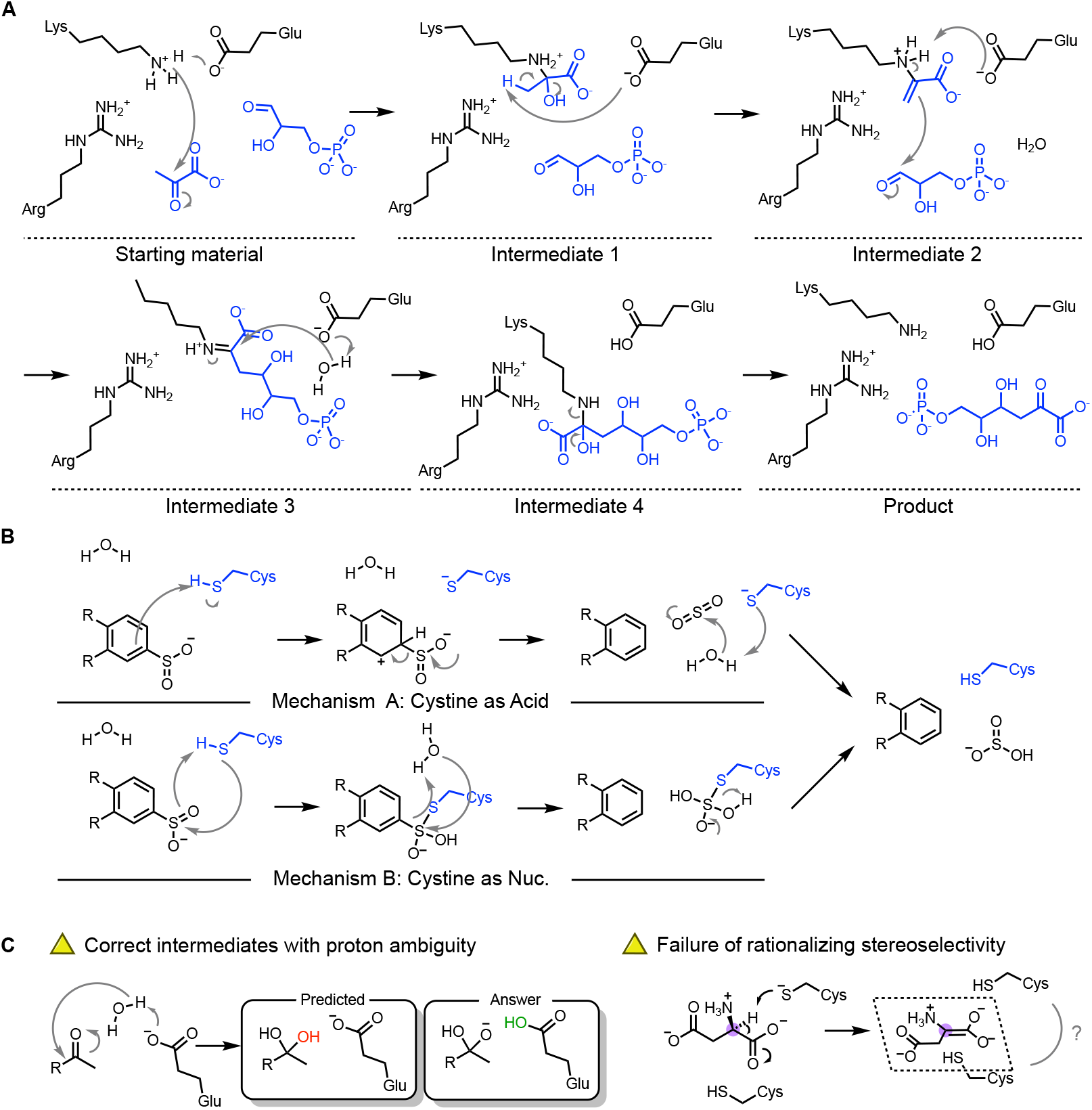
An example of a 5-step ‘Accurate’ mechanism and some limitations of EZSolver. (A) EZSolver correctly chooses lysine to catalyze the aldol reaction. The 5-step mechanism shows the importance of the bidirectional beam search in mitigating error accumulation. (B) Unseen desulfination mechanisms proposed by EZSolver with divergent cysteine functionalities, showcasing its ability to predict insightful enzymatic pathways. (C) Despite assistance from the 2D and chemistry evaluators, EZSolver sometimes fails to predict precise proton distributions, presumably due to the difficulty of curating detailed proton transfer data. While the 2D evaluator provides stereochemical information to guide EZSolver, stereoselectivity cannot be fully rationalized by the model without the exact 3D enzyme environment.

However, despite these advancements, certain fundamental challenges remain. As shown in Figure 6C, the accurate prediction of proton distribution and specific protonation states continues to be a major hurdle. While EZSolver attempts to bridge this gap using the knowledge gained from EZFlow and expert-designed evaluators, predicting a precise hydrogen-bonding network remains exceedingly difficult without explicit 3D conformational data and substrate docking information. Furthermore, similar to its predecessor FlowER, the current EZFlow architecture neglects stereochemical information during the core inference process. Although we deliberately incorporated stereochemical information within the 2D Navigator to focus the model’s attention on corresponding chiral centers, the algorithm still lacks the spatial reasoning required to rationally predict the exact stereoisomer produced within a specific enzyme pocket (Figure 6B). These limitations suggest that while 2D topology and bidirectional graph search can solve the majority of the mechanistic puzzle, the final refinement of stereochemistry and proton-shuttling will likely necessitate the integration of 3D spatial descriptors or hybrid QM/ML approaches in future iterations. By positioning EZSolver as a high-throughput mechanistic generator, these “Plausible” candidates can serve as an optimized starting pool for more computationally expensive 3D-structural or quantum mechanical validations.

### EZSolver outperforms template-based methods on out-of-distribution tasks

To further validate the robustness of this architecture, we benchmarked EZSolver against EZMechanism^42^ and MechFind,^43^ two rule-based predictors. Both models perform excellently on in-distribution tasks, where reaction mechanisms are predicted using a pre-established rule pool curated from the exact same reaction data. EZMechanism incorporates 3D energy calculations to refine searches within its pre-established rule pool, and MechFind utilizes MILP to efficiently retrieve mechanisms. However, their performance remains constrained by the inherent limitations of a fixed dictionary when predicting unseen chemistry. For a rigorous out-of-distribution (OOD) comparison, we adopted the evaluation framework using 55 reactions from the M-CSA dataset used by Ribeiro et al.^42^ for EZMechanism.

While EZMechanism was evaluated by removing only the individual rules for each specific target reaction one at a time, we subjected EZSolver to a much stricter challenge. Specifically, we simultaneously excluded the elementary steps of all 55 target reactions from the training set and trained a single model for this out-of-distribution task. To test MechFind in a similar setting, we only removed rules that were exclusively associated with these 55 reactions; rules shared with other reactions in the remaining training set were intentionally preserved. As shown in Table 1, rule-based models deteriorated significantly in the OOD test. MechFind struggled to generalize without specific templates, yielding a mere 27.3% retrieval rate and an 18.8% accurate rate. On the other hand, despite the more challenging training constraint, EZSolver performed at a comparable level to our previous analysis. By utilizing ensemble sampling across five independent runs, the accuracy reached 52.7%, and the overall mechanism retrieval rate climbed to 72.7%. Furthermore, the fact that the plausible rate remains remarkably close to the retrieval rate underscores the robust chemical intuition embedded in EZSolver. This substantial improvement highlights the core advantage of our generative approach: while deterministic rule-based models are limited by manually curated reaction rules, EZSolver discovers complex and non-standard pathways by effectively allocating computational time to explore the mechanistic landscape.

**Table 1.** Performance comparison of EZsolver to template-based mechanism prediction methods on a strict out-of-distribution (OOD) task. Because EZMechanism lacks open-source checkpoints and training data, its OOD metrics are taken directly from the original publication. Mechanism proposals from MechFind were evaluated under the same categorization as EZSolver. The ‘Ensemble’ method aggregates the outputs from five independent runs of EZSolver, demonstrating that inference-time scaling further boosts the exact accuracy. Overall, the severe performance degradation of deterministic rule-based models on unseen templates highlights their inherent generalization limits, explicitly showcasing the extrapolative advantages of our template-free generative framework.

| Model | Exactly Matched | Accurate | Plausible | Retrieved | Inference time | Model |
| --- | --- | --- | --- | --- | --- | --- |
| EZMechanism | 23.6% | -- | -- | 50.9% | hours | Rule-based |
| MechFind | --- | 18.8% | 25.5% | 27.3% | seconds-minutes | Rule-based |
| **EZSolver** | -- | 36.4% | 54.6% | 61.9% | minutes | Rule-free |
| **EZSolver (Ensemble)** | -- | 52.7% | 70.9% | 72.7% | hours | Rule-free |

## CONCLUSION

Enzymatic reaction mechanism prediction is challenged by limited data, vast reaction spaces involving complex biomolecules, and the presence of reversible reactions. These make deterministic architectures such as sequence-to-sequence or graph-to-sequence models, or even rule-based methods used in organic reaction prediction highly susceptible to hallucinations and unable to extrapolate to out-of-distribution data. To address these intrinsic limitations, we introduce EZSolver, a bidirectional searching model powered by a flow-matching-based single-step predictor, EZFlow. EZFlow is built on the architecture of FlowER and is finetuned to capture the catalytic functionalities of amino acid residues within enzymatic active sites. Most importantly, by incorporating inverse flow, EZFlow overcomes the single-directional inference constraint and becomes capable of predicting elementary steps across the entire thermodynamic landscape, resolving the issues caused by reversible reaction pairs commonly found in enzymes.

For single-step predictions, benchmarking demonstrates that the intrinsic bias of deterministic models poses a significant hurdle in predicting enzymatic elementary steps. Conversely, our model successfully predicts not only the ground truth (72.4% accuracy) but also a broad manifold of chemically plausible elementary steps. For entire mechanism inference, the evaluator-guided bidirectional beam search algorithm, which integrates expert knowledge to focus the search on correct chemical tracks, proposed 84.6% chemically plausible and 60.0% accurate chemical mechanisms, with an average inference time of 1500 seconds per reaction.

Furthermore, our out-of-distribution benchmarks against rule-based models underscore the distinct advantages of this generative framework. While rule-based systems are strictly capped by the boundaries of manually curated dictionaries of rules, EZSolver bypasses these rigid template restrictions, learning the fundamental chemistry of electron redistribution to generate novel elementary steps. As the generative framework tailored for enzymatic pathways, EZSolver opens the room for flexible and chemistry-informed predictions. Although challenges remain regarding precise 3D stereochemistry and the inherent data ambiguities of protonation states, EZSolver provides a robust chemical foundation that could integrate with computationally intensive QM/MM or 3D structural refinements. Additionally, through careful curation of transition metal-catalyzed organic reaction data, future training could empower the model to decipher complex metalloenzyme mechanisms, such as heme catalysis. Ultimately, this could enable EZSolver to expand its applicability across the entire enzymatic landscape. We believe this template-free approach represents a crucial step toward predictive enzymology and the discovery and design of novel biocatalysts for sustainable chemistry.

## ASSOCIATED CONTENT

### Supporting Information

Code and data, details of data curation and the similarity analysis, discussion of inverse flow, and further experimental and computation results.

### Data Availability Statement

All code and data used to produce the results in this study are provided at https://github.com/fhalab/EZSolver.

## AUTHOR INFORMATION

Notes

The authors declare no competing financial interest.

## ACKNOWLEDGMENT

This work was supported by the National Science Foundation Division of Molecular and Cellular Biosciences (MCB-2016137 to FHA). J.Y. was supported by an NSF Graduate Research Fellowship and the Google PhD Fellowship.

