## Supporting Information for "EZSolver: Template-free prediction of polar enzymatic mechanisms via bidirectional flow matching and search"

### Contents


**1 Data curation** **S3**

**2 Inverse flow** **S5**

**3 Algorithm of bidirectional beam search** **S7**

**4 Hyperparameters** **S8**

**5 Benchmark of EZFlow and EZSolver** **S8**

**6 Evaluators in EZSolver** **S11**

### 1 Data curation

The M-CSA database[1] has around 1000 detailed mechanisms for enzymatic reactions. These mechanisms are represented as reactants, all the intermediates, and products, across all elementary steps. To curate the data, the corresponding Marvin files of each elementary step were first downloaded. Subsequently, the structures in Marvin files were converted to SMILES using RDKit. We realized that chemical structures might suddenly appear or disappear in the middle of reactions, where in most cases, are products or additional starting materials. This causes an imbalance of atoms within elementary steps. To address the problem, reactants and products of overall reactions are fitted into the imbalanced reactions to meet atom and charge balance. Ultimately, we retrieve around 2000 elementary steps. Next, reactions having radicals or metals are removed, and heavy atoms in reactants and products are mapped using RXNmapper. The hydrogens on each heavy atom are compared between reactants and products. If the count remains the same, those hydrogens are mapped directly. If the counts differ, the heavy atom that loses a proton is paired with the heavy atom that gains one. Finally, approximately 1200 elementary steps are curated for training and testing.

A subset of 186 elementary steps, which consists of 65 full reaction mechanisms, was selected for testing model performance in most results presented in this study. Crucially, not a single elementary step in this testing set shares an identical SMILES string with the training data. Further analysis of the Reaction Tanimoto Similarity between elementary steps from the training and testing sets revealed that roughly 13% (25 steps) of the testing data exhibited a 100% maximum reaction similarity compared to the training data (Figure S1A). A detailed breakdown showed that 22 of these steps are proton transfers between catalytic residues. The remaining 3 steps involve ester hydrolyses sharing the same substructures with the training data. Similarly, in the EZSolver benchmark study, zero steps in the testing set share identical SMILES strings with the benchmark training data. Within this specific testing set, 19 out of 105 elementary steps exhibit a 100% reaction similarity. Of these, 15 are proton transfers between residues, while the remaining 4 steps consist of sugar tautomerizations and an amide hydrolysis.

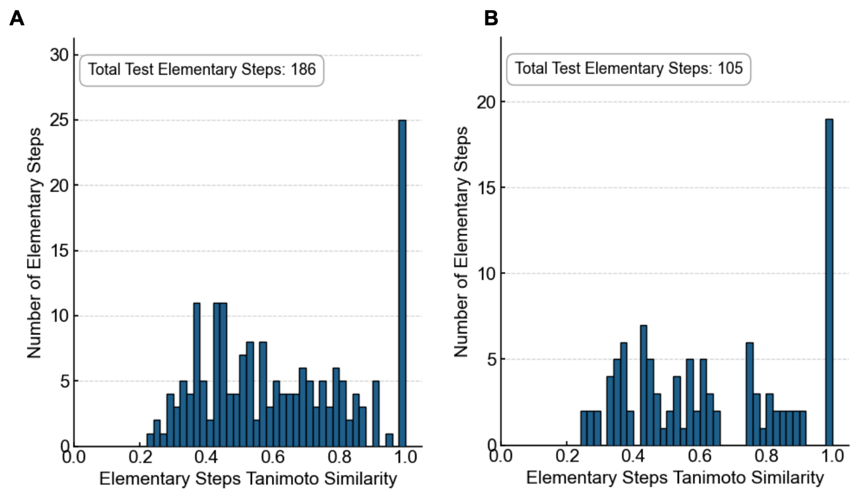


Figure S1: **Distributions of Reaction Tanimoto Similarity between the training and testing sets.** (A) The dataset split used for evaluating EZSolver. (B) The benchmark dataset split used for comparing EZSolver against MechFind and EZMechanism.

### 2 Inverse flow

FlowER[2] is a diffusion model that operates on the Bond-Electron (BE) matrix and is conditioned solely on the atom types within given compounds. This architecture makes it fundamentally impossible to learn reversible reaction pairs. Because the objective flows for forward and reverse reactions are exactly opposite, yet operate under identical conditioning, the model would receive contradictory training signals if both were present in the same dataset. This is further proven by overfitting experiments of two elementary steps that failed when they were the reverse of each other. Coley and co-workers circumvented this limitation by rigorously curating their training data using expert-derived templates and hard-coding pKa values, thereby restricting the reaction space to encompass unidirectional transformations. While this curation prevents conflicts of objective flow during training, it essentially forces the model to ignore the reversibility inherent in real chemical systems. Consequently, FlowER functions strictly as a unidirectional reaction predictor, rendering it inadequate for modeling the nuanced, reversible steps essential to comprehensive enzymatic mechanisms even after decent finetuning.

Recognizing that the model is conditioned solely on atom types, which remain invariant between the reactant and product states, we hypothesized that the flow-matching ODE could be reconceptualized not merely as a generative trajectory, but as a bidirectional thermodynamic vector. By reversing the time integration during inference, the sample distribution would be guided from the ’product’ state backward toward the ’reactant’ state. This ‘inverse flow’ approach theoretically enables the prediction of elementary steps that are oppositely aligned with the training set.

To validate this hypothesis, we used MMFF to roughly estimate the thermodynamic profile of each elementary step and segregated the dataset into exothermic and endothermic subsets (Figure S2A). Analysis of Reaction Tanimoto Similarity between endo-subset and exo-subset is shown in Figure S2B. By applying inverse flow, a model preliminarily finetuned solely on the exothermic dataset achieved a 44% prediction accuracy when applying inverse flow to the unseen endothermic dataset (Figure S2A). This extrapolation confirms that implementing inverse flow successfully empowers the architecture to extrapolate across opposite thermodynamic profiles. Additionally, by defining the reaction ’direction’ alongside the thermodynamic profile, we effectively resolve the contradictory training signals.

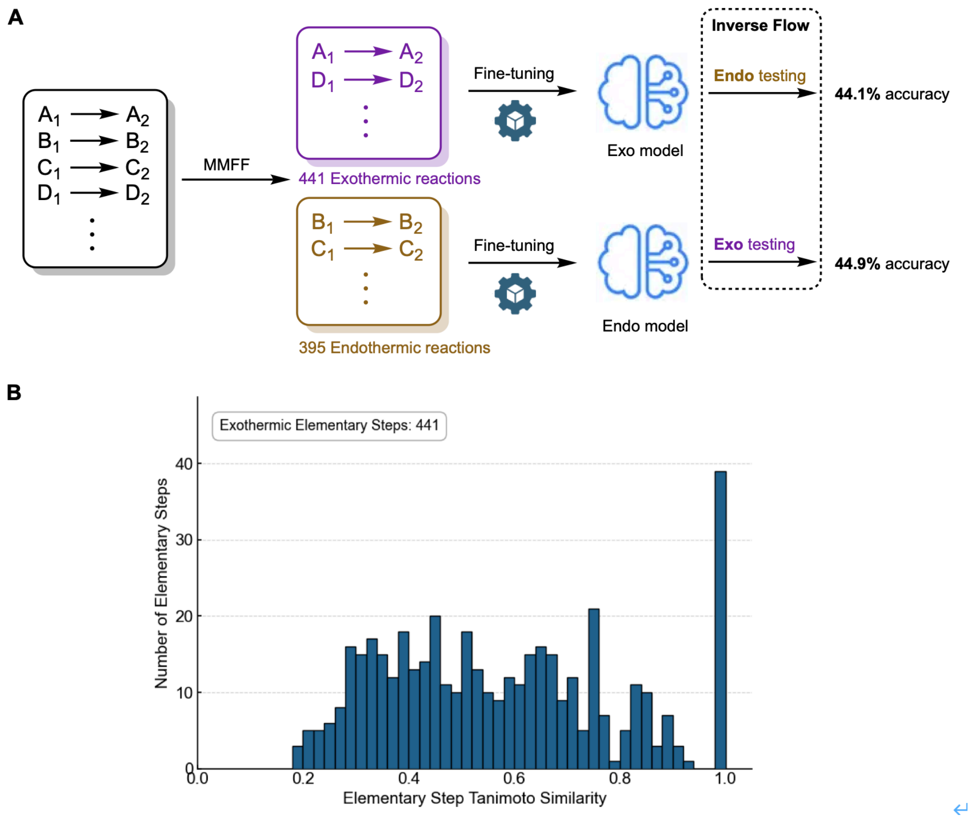


Figure S2: **Inverse flow enables robust extrapolation across reversed thermodynamic landscapes.** (A) Reactions were segregated into exothermic and endothermic subsets based on MMFF estimations. Preliminary finetuning on each datasets yield an Exo model and an Endo model. Cross inference of these two models on the thermodynamically opposite dataset through inverse flow gave decent accuracy. (B) Reaction Tanimoto Similarity of Exothermic reactions on Endothermic reactions.

### 3 Algorithm of bidirectional beam search

**Algorithm 1** EZSolver: Dual-Evaluator Guided Bidirectional Beam Search

**Input:** Reactant $R$, Product $P$, Generative Model $\mathscr{M}$, Max depth $D$, Beam width $W$
**Initialize:** Active beams $\mathscr{X}_{fwd} \leftarrow [R]$, $\mathscr{X}_{bwd} \leftarrow [P]$
**Initialize:** Visited states $\mathscr{H}_{fwd} \leftarrow \{R\}$, $\mathscr{H}_{bwd} \leftarrow \{P\}$
**Initialize:** Valid pathways $\Omega \leftarrow \emptyset$
**for** $d = 1$ **to** $D$ **do**
    **if** $\mathscr{X}_{fwd}$ and $\mathscr{X}_{bwd}$ are both empty **then**
        **break**
    **end if**
    **for each** direction **in** $\{\text{forward}, \text{backward}\}$ **do**
        Generate and score next states from the active beam using $\mathscr{M}$ ▷ Filtered by 2D Navigator & Chem Evaluator
        Discard states already present in the current direction's visited set $\mathscr{H}$ ▷ Graph loop prevention
        $Candidates \leftarrow []$
        **for each** valid new state $S$ **do**
            **if** $S$ exists in the opposite direction's visited set **or** $S$ is the target **then**
                Construct the full reaction pathway and add to $\Omega$
            **else**
                Append $S$ to $Candidates$
            **end if**
        **end for**
        Update active beam: Select top $W$ scoring states from $Candidates$
        Add selected states to the current direction's visited set $\mathscr{H}$
    **end for**
**end for**
**Return:** $\Omega$ sorted by pathway length and average score

### 4 Hyperparameters

Table S1: Hyperparameter setting used in the experiments. Best hyperparameters are highlighted in **bold**.

| Model | Task | Parameters | Values |
|---|---|---|---|
| EZFlow | Training | Attention encoder layers | 12 |
| | | Attention encoder heads | 32 |
| | | Sigma | **0.05**, 0.1, 0.15 |
| | | Learning rate | **0.0001**, 0.00001 |
| | | Scheduler | NoamLR |
| | | Embedding size | 256 |
| | | Hidden size | 256 |
| | | RBF low, high, step | (0, 12, 0.1) |
| | | ODE solver method | Dopri5 |
| | | ODE solver absolute error tolerance | 0.0001 |
| | | ODE solver relative error tolerance | 0.0001 |
| EZFlow | Inferencing | Sigma | 0.05, **0.08**, 0.1 |
| | | Sample size | 50, 100, **200** |
| EZSolver | Inferencing | Beam width | 5, **10**,15 |
| | | Max beam depth | 3, **4**, 5 |

### 5 Benchmark of EZFlow and EZSolver

#### EZFlow

In the benchmark experiment against FlowER,[2] Chemformer,[3] and Graph2SMILES,[4] all the models were finetuned on *train.txt* based on the suggested protocols. With prior knowledge, we adapted focal weights in finetuning FlowER and tested it with the optimal sigma. The results are shown in Table S2.

| Models | top-1 | top-5 | top-10 | top-30 | top-50 | top-100 | top-200 |
|---|---|---|---|---|---|---|---|
| EZFlow | 9 | 25 | 41 | 54 | 58 | 66 | 72 |
| FlowER | 4 | 15 | 21 | 34 | 37 | 40 | 44 |
| Chemformer | 8 | 11 | 13 | 18 | 20 | 21 | 25 |
| Graph2SMILES | 0 | 0 | 0 | 0 | 0 | 0 | 0 |

Table S2: Benchmark results on *test.txt*

Chemformer suffers from severe hallucinations in enzymatic elementary step prediction. Figure S3 shows the top-4 predictions from Chemformer on an elementary step of serotonin acetylation. All the predicted intermediates feature new motifs along with the absence of serotonin substructures. Detailed investigation of the top-200 predicted intermediates showed that none of them obeys the conservation of atoms.

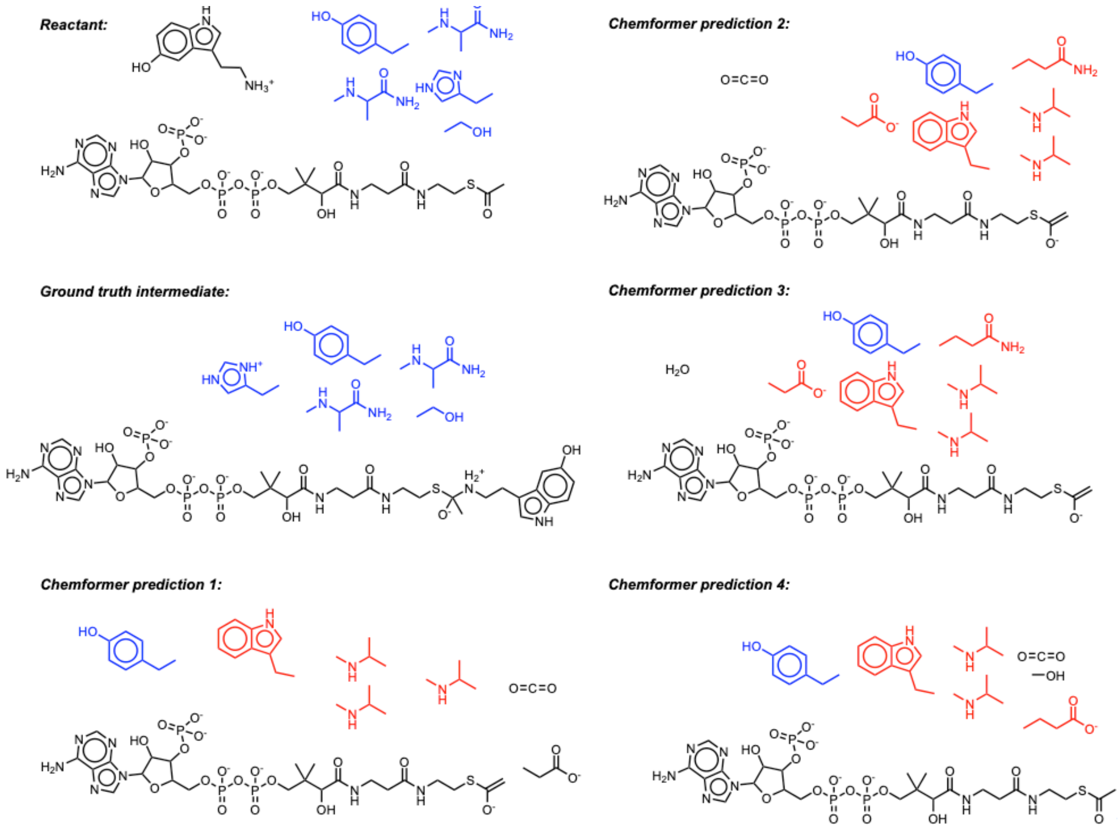


Figure S3: Examples of the severe hallucinations from Chemformer on enzymatic elementary step predictions.

#### EZSolver

Because the checkpoint files for EZMechanism[5] are not publicly available, we conducted the benchmarking evaluation strictly on the same data split utilized in their out-of-distribution (OOD) assessment. By removing reaction rules that were uniquely derived from each testing reaction, EZMechanism successfully retrieved 28 out of 55 reaction mechanisms, among which 13 aligned identically with the ground truth. Notably, this evaluation was performed independently across the 55 reactions, necessitating the removal of specific rules on a reaction-by-reaction basis.

In contrast, rather than training 55 independent models, we demonstrated the robust generalization of EZSolver by training a single unified model; this was achieved by removing all elementary steps associated with these 55 reactions entirely from the training dataset. For MechFind,[6] the unique rules originating from these 55 reactions were likewise removed. Finally, predicted trajectories from both EZSolver and MechFind were rigorously reviewed and categorized into three distinct fidelity levels: 'Accurate', 'Chemically Plausible', and 'Retrieved'.

While PMechRP[7] is a relevant state-of-the-art model for mechanism prediction of organic reactions, its unique hybrid architecture restricts its adaptability to polar enzymatic reactions. Because standard finetuning protocols cannot be applied symmetrically to this framework, a direct baseline comparison would be unfair. Therefore, PMechRP was omitted from the benchmarking discussion to ensure a fair evaluation.

### 6 Evaluators in EZSolver

#### 2D Navigator

Biomolecules have complicated structures capable of undergoing a vast array of chemical transformations. Consequently, while EZFlow predicts a multitude of potential intermediates, most of these do not align with the chemical pathway toward the final product. To address this, the 2D Navigator is designed to selectively filter and prioritize intermediates that undergo electron redistribution (bond formation or cleavage) specifically on relevant reactive atoms. By comparing the Bond-Electron (BE) matrix of the current state with that of the final product state, the changes in electron counts on these reactive centers are tracked, enabling the generation of a dynamic 2D reactive atom mask.

Furthermore, recognizing that neighboring atoms adjacent to the reaction core may participate in the mechanism, such as pericyclic reactions, the 2D mask is spatially extended. The predicted intermediates are subsequently ranked based on a reward function derived from this extended 2D mask, providing an efficient structural screening prior to downstream evaluation by the Chemistry Evaluator.

#### Chemistry Evaluator Rules and Penalty Metrics

To suppress non-physical generative hallucinations and steer the bidirectional beam search toward chemically legitimate mechanisms, we implemented a rule-based expert system called the Chemistry Evaluator. This system screens elementary step candidates on an atom-by-atom basis across the core reaction landscape, imposing structural rewards (**negative costs**) or geometric penalties up to infinite cost (forbidden pathways). The detailed criteria categorized by atom type are elaborated below:

##### Enzymatic Amino Acid Protection

To preserve the structural and chemical integrity of the enzyme active site, specific filters are applied to atoms flagged as part of catalytic amino acid residues:

- **Carbon Skeleton:** For any carbon atom belonging to an amino acid side chain (e.g., Asp, Glu), its neighboring atom composition and hybridization state ($sp^2$ versus $sp^3$) must remain completely unchanged. Any discrepancy triggers an infinite penalty ($\infty$), effectively preventing irrelevant orthoacid formations and side-chain transformations.

- **Resonance Nitrogen Stabilization:** Inactive nitrogen atoms within highly resonance-stabilized functionalities, such as the guanidinium group of Arg or the amide groups of Asn and Gln, are forbidden from acting as nucleophiles. If an un-aromatic nitrogen is adjacent to an $sp^2$-hybridized carbon, any change in its heavy-atom coordination degree is strictly blocked ($\infty$).

**Carbon Chemistry Rules**

Carbon atoms undergo rigorous screening to evaluate proper oxidation states, hybridization, and transient chemical intermediates:

- **Oxocarbenium Intermediate Reward:** The formation of high-energy yet highly favorable transient oxocarbenium (oxonium) ions is actively incentivized. If a carbon transitions from a neutral state into either a positively charged state conjugated with oxygen ($[C^+]$–O) or a double-bonded oxonium moiety (C=$[O^+]$), a reward ($\Delta$cost $= -200.0$) is bestowed.

- **Hybridization and Substitution Constraints:** Unrealistic $sp^2 \rightarrow sp$ conversions (e.g., generating highly unstable allenes or ketenes in enzymatic channels) are entirely penalized ($\infty$).

- **Prohibition of $sp^2$ Direct Substitution:** Direct nucleophilic substitution on an $sp^2$-hybridized carbon framework (such as carbonyl positions) without proceeding through a tetrahedral intermediate is forbidden ($\infty$).

- **Anomeric $sp^3$ Substitution Screening:** For geometrically allowed $sp^3 \rightarrow sp^3$ substitution pathways (e.g., glycosidic bond cleavage via $S_N1$ or $S_N2$ mechanisms), the evaluator applies hierarchical scoring based on the carbon's oxidation state. Substitutions occurring at oxygen-dense anomeric or acetal centers (oxidation state OS $\geq 2$) receive a substantial reward ($\Delta$cost $= -100.0$), whereas unfavored direct substitutions on unactivated aliphatic carbons are penalized ($\Delta$cost $= +100.0$).

**Phosphorus Chemistry Rules**

For phosphorus atoms—predominantly governing kinase, phosphorylase, and phosphatase reactions—the evaluator monitors coordination geometry:

- **Associative and Dissociative Trajectories:** Transitions between a 4-coordinate tetrahedral phosphate and a 5-coordinate trigonal bipyramidal intermediate are rewarded ($\Delta\text{cost} = -50.0$) to facilitate phosphate transfers.
- **Metaphosphate or Coordination Protection:** For 4-coordinate phosphorus frameworks surrounded by at least three oxygen atoms, any substitution that alters the identity of the heavy-atom neighbors without going through a proper 5-coordinate intermediate is heavily restricted ($\infty$).

##### General Topological and Ring Strain Rules

Beyond specific atom identities, a global Smallest Set of Smallest Rings (SSSR) analysis is performed to assess thermodynamic and kinetic feasibility against ring strain and entropic penalties:

- **Macrocycle Forbidding:** The formation of new large rings ($\geq$ 8-membered cycles) is assigned an infinite cost ($\infty$) due to severe entropic penalties in standard enzymatic pockets.
- **Geometric Ring Strain:** The creation of highly strained 3- or 4-membered rings (such as epoxides or aziridines) is tightly controlled. If the newly stitched small ring contains any $sp^2$ or sp hybridized atoms, the prohibitive angle strain results in an immediate infinite penalty ($\infty$). Conversely, all-$sp^3$ small rings are permitted but heavily penalized ($\Delta\text{cost} = +1000.0$), ensuring they are only selected when open-chain pathways are structurally impossible.
